# Sequential sensory integration drives foraging decisions in leaf-cutting ants: Volatiles, contact cues and phytochemistry

**DOI:** 10.64898/2026.05.29.725965

**Authors:** Ayelén Nally, M. Soledad Méndez, Patricia Carina Fernández, Fernando Federico Locatelli

## Abstract

Identifying the sensory cues that enable insects to find host plants, and understanding the neurobiology underlying their selection, provide solid foundations for developing state-of-the-art pest management strategies. Our work was aimed at identifying the main sensory cues attracting the leaf-cutting ant *Acromyrmex ambiguus* to alternative host plants in commercial willow plantations in the Lower Delta of the Parana River (Argentina), with a focus on native plant species. Eight plant species were selected and evaluated as potential hosts under field and laboratory conditions, allowing the establishment of a robust hierarchy of preference based on individual and collective behaviour. As a result, *Senna corymbosa* emerged as the most preferred species, whereas *Blepharocalyx salicifolius* was the least preferred. Video analyses of ant foraging in controlled indoor nests revealed a sequential decision-making process underlying plant preference and consumption. This included an initial approach driven by olfactory cues, followed by a second step involving contact-dependent cues that elicited leaf-cutting and carrying the leaf fragments to the nest. Volatile compounds and leaf cuticular components potentially involved in plant preference were identified. In addition, physicochemical analysis of both plant species - including total sugars, organic matter, polyphenols, leaf hardness, total proteins and lignin-revealed differences, particularly in polyphenol content, which may contribute to preference patterns. These findings provide insights into the sensory ecology of host preference and inform management strategies based on the reintroduction of native plants as alternative resources in willow plantations, potentially reducing pesticide use and promoting environmental sustainability.

**Summary statement:** This work shows how leaf-cutting-ants rely on olfactory and contact cues, sequentially, for foraging decision-making of plant species. Thus, revealing how sensory cues shape their foraging decisions.

## Introducción

Leaf-cutter ants (Order: Hymenoptera; Family: Formicidae; Subfamily: Myrmicinae; Tribe: Attini) are among the invertebrates that cause the greatest damage to plantations in the Neotropics (Fowler, 1978). This damage results from the collection of plant fragments, which are transported to underground nests where the material is used to cultivate a symbiotic fungus that serves as the sole food source for the larvae (Littledyke and Cherrett, 1976; Hölldobler and Wilson, 1990; Fowler and Robinson, 1979; Forti et al., 2006).

The effects of defoliation include a reduction in photosynthetic area, decreased energy allocation to growth, increased susceptibility to pathogens due to tissue damage and reduced energetic reserves, and delayed leaf production, which further increases plant vulnerability (Rockwood, 1973). The damage is particularly detrimental during the first three years of a tree’s life (dos Santos et al., 2013). As a consequence, leaf-cutter ants are considered the dominant herbivores in the Neotropics, the most important agricultural pests in the Americas (Cherrett, 1986; Fowler, 1978), and among the most economically significant pests in South America (Scherf et al., 2022; Mikheyev, 2008; Erthal et al., 2004).

Plant species or varieties cultivated in forest plantations are often selected for high growth rates at the expense of defensive traits (Jiménez, 2022). Moreover, exotic commercial species such as *Salix babylonica* var. *Sacramenta*, which represents nearly 60% of the commercial forest plantations in the study region (Argentine Ministry of Economy, 2023), have not coevolved with leaf-cutter ants, suggesting a lack of specific defenses against these herbivores. Additionally, leaf-cutter ants prefer plant material with fewer physicochemical defenses, generally associated with young leaves (Vasconcelos & Cherrett 1995; Farji-Brener, 2001). It is well documented that leaf-cutter ants Acromyrmex spp. reduce wood production and affect tree establishment in forest plantations (Montoya-Lerma et al., 2012). Their effect can be as much as a reduction of up to 32% in height, 25% in circumference, and 60% in wood production (Della-Lucía, 1993). Particularly, *Acromyrmex ambiguus* (Emery, 1888) is one of the most important predators of young forest plantations (Sanchez-Restrepo et al., 2019) with an annual impact on dry material of 26 kg/ha in Lower Delta Forestal production area (Jimenez, 2022).

Consequently, the control of leaf-cutter ant populations is particularly intensive during the first months of a tree’s life, when seedlings are most vulnerable, and throughout the first two years after planting (Lewis and Norton, 1973; Vasconcelos and Cherrett, 1997; Nickele et al., 2012). These control measures may account for up to 75% of the pest management budget (Cantarelli et al., 2008), as economic losses caused by leaf-cutter ants are estimated at millions of US dollars annually (Bacci et al., 2009).

In contrast, sustainable agriculture aims to reduce insecticide use while considering the ecological role of ants in soil productivity. However, there is still a lack of information regarding native plant species in forest agroecosystems, which hinders the design of sustainable management strategies (Della-Lucia, 2003). Currently, the landscape is undergoing continuous anthropogenic modification, resulting in only remnants of the native forest remaining (Fracassi, 2017; Kalesnik et al., 2008). These remnants represent approximately 4% of the local vegetation in less frequently flooded areas, whereas more than 80% of the surface is covered by herbaceous plants (Kandus, 2019; Salvia et al., 2010; Morandeira, 2017).

This contrasts with the well-established role of biodiversity in maintaining the functioning of agroecosystems (Altieri et al., 1999; Swift et al., 2004). In the present study, we focused on the push–pull strategy, which combines a “push” stimulus that repels pests with a “pull” stimulus that attracts them (Cook et al., 2007). Previous studies have shown that push–pull strategies can successfully reduce herbivory pressure by leaf-cutting ants in forest plantations through the use of spontaneous vegetation and farnesol (Perri, 2020). However, there is still no information regarding the use of native plant species and the semiochemicals that may act as push or pull stimuli in this particular region.

This study aims to understand behavioral and phytochemical mechanisms underlying foraging decisions in *A. ambiguus*. We hypothesize that leaf-cutting ants have the capacity to detect and identify the preferred and rejected vegetal species based on phytochemical cues. Furthermore, we hypothesize that some native species from the region are more preferred than the commercial willow species, whereas others are less preferred. To test these hypotheses, we evaluated sensory modalities involved in plant recognition and characterized the behavioral responses to the most and least collected plant species. Finally, we identify native species with potential applications in sustainable pest-management strategies.

## Materials and methods

### Biological material

*A. ambiguus* colonies, including symbiotic fungus (*Leucocoprinus gongylophorus*), were collected at the Delta Agricultural Experimental Station of the National Institute of Agropecuary Technology (INTA EEA Delta) Campana, Buenos Aires, Argentina during 2017 and 2019. Each colony was collected and placed in plastic boxes (32 x 42 x 6.5 cm). A larger box would be analogous to the general structure of the colony, and smaller rectangular boxes (23 x 14 x 7 cm) would be analogous to the fungus and brood chambers. Colonies were maintained in a chamber at 23–25°C, 60% r.h., and 12h light:12h dark photoperiod (L12:D12) (Perri et al., 2017).

The colonies were fed daily with plant species not used in the experiments, such as ash leaves fraxinus (*Fraxinus sp.*), bridal wreath spirea (*Spiraea cantoniensis*), tipa (*Tipuana tipu*), jacaranda (*Jacaranda mimosifolia*), and blackberry (*Morus nigra*), among others. They were also given fruits such as apple (*Malus domestica*), orange (*Citrus × sinensis*), and mandarin orange (*Citrus reticulata*). Additionally constant food throughout their stay in the laboratory from their collection, including during the experimental periods, was a mixture of oat flakes, maize flour, and rice. Behavioral experiments were conducted 3 months after the colony was collected, in order to ensure the turnover of the forager ants in the colony, that is, that these ants had no previous experience with the plant species used in the experiments. All experiments were carried out in the spring-summer period.

Native plant species used in lab experiments were *Blepharocalyx salicifolius*, *Myrsine laetevirens*, *Terminalia australis*, *Erythrina crista-galli*, *Sesbania punicea*, *Senna corymbosa*, *Verbena bonariensis*. All of them were kept in pots under natural conditions in the greenhouse of the Institute of Physiology, Molecular Biology and Neurosciences. Trees used in field experiments were located at INTA EEA Delta (−34.17 lat, −58.86 long) at patches of native flora with composition and location of specimens differing between patches. *B. salicifolius*, *M. laetevirens*, *E. crista-galli*, and *S. corymbosa* were present on the three patches. The patches were located on both banks of a stream adjacent to a poplar and willow plantations.

### Herbivory in the field

Each specimen within the patch was photographed against a gray background once a month from January to April 2019 following Perri’s (2020) methodology. This period coincides with ant’s peak activity. Photographs were taken for each specimen to assess their foliage density. The first photograph was taken as 100% of the foliage. Subsequent photographs were compared with the first to estimate a % of foliage lost. If due to growth, photographs showed more foliage than the initial one then the new measurements were considered as 100%.

### Herbivory in the laboratory

Dual choice tests were performed by evaluating the following species: the commercial willow (*S. babylonica* var. Sacramenta) -one of the most widely used in forest plantations in the area - and 7 native plants maintained in IFIBYNE Institute. During each trial, the ants were able to cut and collect portions of the fresh leaves offered to them. Each trial lasted 45 minutes. The leaves were photographed before and after the trial (Fig. 2.B) and then their area was quantified by ImageJ (Guerrero Rincón et al., 2012). The percentage of surface consumed for each leaf was calculated, and the difference in consumption between the two leaves was calculated as the difference between the respective percentages. Each trial consisted of the presentation of a pair of leaves. The eight plant species used were presented in pairs. This generated 28 possible combinations. In addition, control trials were conducted with two leaves of the same species to establish a reference value indicating the expected difference in consumption between two leaves with identical characteristics. In total, each repetition of the experiment included 36 trials that were conducted with 6 colonies of *A. ambiguus*. This generated a total of 216 trials. In all cases where plant leaves were used, they were of the same maturity. Experimental setup: Each ant colony was connected to an experimental arena via a wooden bridge. Two fresh leaves of similar size from the plant species used were placed in the experimental arena (Dimensions: 43 cm x 11 cm x 6 cm). The experimental arena was filmed with a webcam (Logitech C920 Full 30 FPS) from the moment an ant made physical contact with the experimental arena until 45 minutes afterward (Fig. 2.A).

**Figure. 1.**
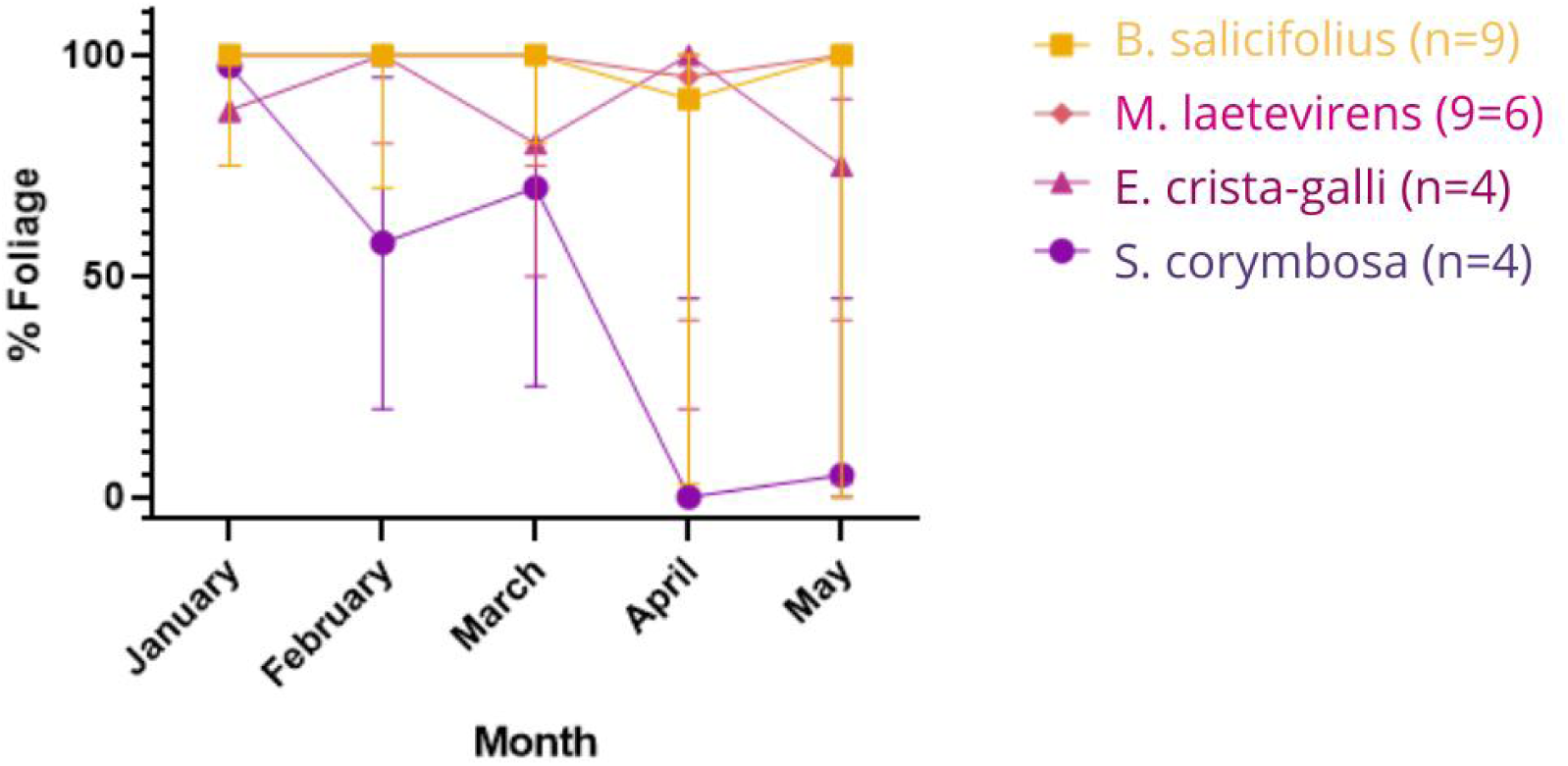
Percentage of foliage of native plant species over time in Summer-Autumn 2019 in Lower Delta of the Parana river, Buenos Aires, Argentina. Medians and 95% confidence intervals are shown. *B. salicifolius,* (N=9); *M. laetevirens*, (N=6); *E. crista-galli*, (N=4); and *S. corymbosa*, (N=4).

**Figure 2.**
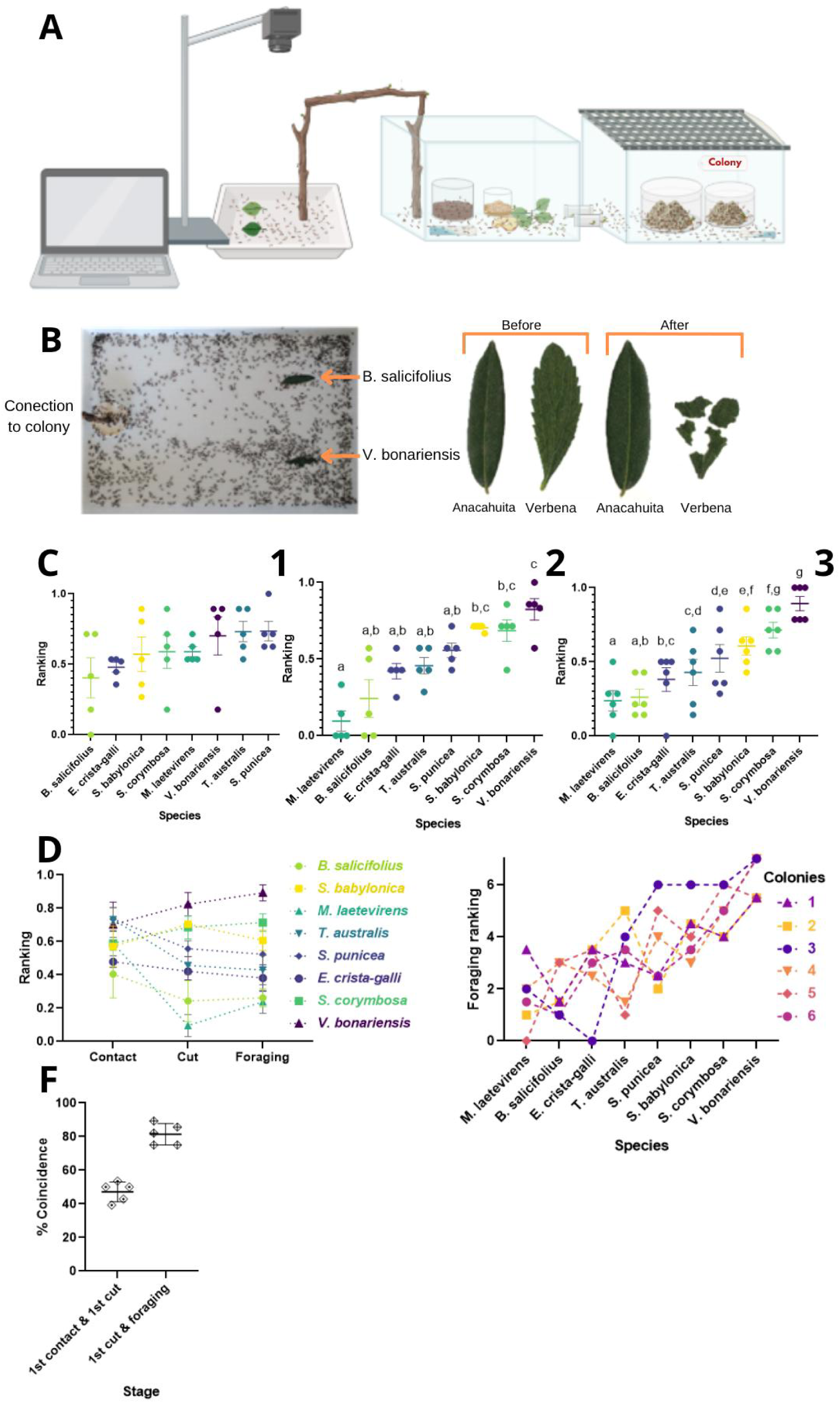
Consumption hierarchy in the laboratory. A. Experimental device for dual choice tests with fresh plant leaves. B. Left panel: Presentation of fresh leaves of the native species *B. salicifolius* (top) and *V. bonariensis* (bottom) in a dual consumption test. Right panel: *B. salicifolius* (left) and *V. bonariensis* (right) leave before and after the choice test. C. Leaf consumption hierarchy of the study species. Each point represents the proportion of time in which the species indicated on the X axis was the most consumed in a pair preference test for a colony. The horizontal bars and error bars indicate the mean and standard error among colonies. Hierarchy of approach to leaves of the study species where each point indicates the proportion of times in which the plant indicated on the X axis was: 1. First approach of an ant of one of 5 colonies. There were no significant differences between species (Kruskal-Wallis 10.40; p=0.167). 2. First cut of an ant of one of 5 colonies. Different letters indicate significant differences (Kruskal-Wallis 28.89; p<0.01; Dunn contrasts; p<0.05). 3. Consumption of an ant of one of 6 colonies. Different letters indicate significant differences between species (p<0.0001; post hoc comparisons by Kruskal-Wallis and Dunn; different letters indicate p<0.001). D. Hierarchy of contact, first cut, and consumption for each of the species under study. Each point in contact and first cut indicates the mean and standard error of 5 colonies. Each point for consumption is the average of 6 ant colonies. E. Hierarchy of consumption per colony for each of the species under study. F. Percentage of coincidence between: (i) the first physical contact of the foraging ant with a leaf and the first cut made on a leaf, and (ii) between the first cut made on a leaf and the final consumption of the leaves.

A sequence of three instances were identified: 1) 1st approach, the first species touched by an ant; 2) 1st cut, first leaf species cut ; and 3) consumption, leaf species most cut ant the end of the trial. A species hierarchy was calculated for each instance. Approach hierarchy: the leaf with which a forager ant made its first physical contact was identified in each dual trial. In that trial, the chosen species received a score of 1 and the other 0. This was repeated for the 28 leaf pairs. Each species participated in a total of 7 trials. At the end, an index proportional to the number of trials in which it was the first choice was calculated. An index of 1 corresponds to the species being chosen all 7 times, and an index of 0 (zero) corresponds to the species not being chosen at all. Finally, an average of the 6 colonies evaluated was calculated. The same procedure was followed for the first-cut hierarchy. For the consumption hierarchy the procedure was slightly different. The difference in consumption measured in trials conducted with two leaves of the same species (control trials) was taken into account. The average consumption difference for all species and all colonies was used as a reference value—11.87—to evaluate the results of the dual trials (Fig. S1). If the consumption difference between two leaves was greater in absolute terms than the reference value, then a “1” was assigned to the most consumed species and a “0” to the least consumed species. If the consumption difference between the two leaves was equal to or less than the reference value, a tie was considered, and a “0.5” was assigned to both species (Perri et al. 2020). The value of 0.5 for each represents the fact that both were chosen equally. Based on these criteria, each cell of a matrix was filled with 1, 0, or 0.5. The scores were then summed along each row. If a species consistently proved to be the most consumed, it ended the experiment with a score of 7. If, on the other hand, it consistently proved to be the least consumed, it ended with a score of 0. If there was a tie across all trials, it ended with a score of 3.5. To calculate the final consumption hierarchy, the scores from the six colonies were averaged.

### Collective preference assays between least preferred (*B. salicifolius*) and most preferred (*S. corymbosa*) leaf discs

Leaves were collected and immediately cut with a paper hole punch. The size of the discs was controlled by measuring both the hole punch diameter and the disc diameter with a digital caliber (Parkside). Each side of the hole punch was used for only one of the species to avoid contaminations. The leaf discs could be differentiated by the experimenter based on their position in the experimental arena and their abaxial veins.

To assess the collective preference of a leaf-cutting ant colony, a group of 12 fresh leaf discs was presented: 6 *B. salicifolius* discs and 6 Sen discs from the field (Fig. 3.A). These discs were placed intermingled and formed a semicircle, so that all the discs were at the same distance from the bridge that connected the experimental arena to the colony. Between 5 and 3 trials were conducted with each colony. We recorded the time it took for an ant to collect a leaf disc and the species of the collected disc. The ant that collected the disc and the surrounding ants were picked up with forceps and transferred to a holding arena so that the next collection event during the same trial would not be influenced by this behavior. At the end of the experiment, all ants were returned to the colony. None of the discs collected during the experiment reached the colony, so that outgoing foragers would have no information about offered material. Furthermore, removing ants that had picked up a disc from the experiment prevented potential path-marking behavior with pheromones that generate positive reinforcement of the path to the collection site and also other possible odor traces from the collected leaf.

**Figure 3.**
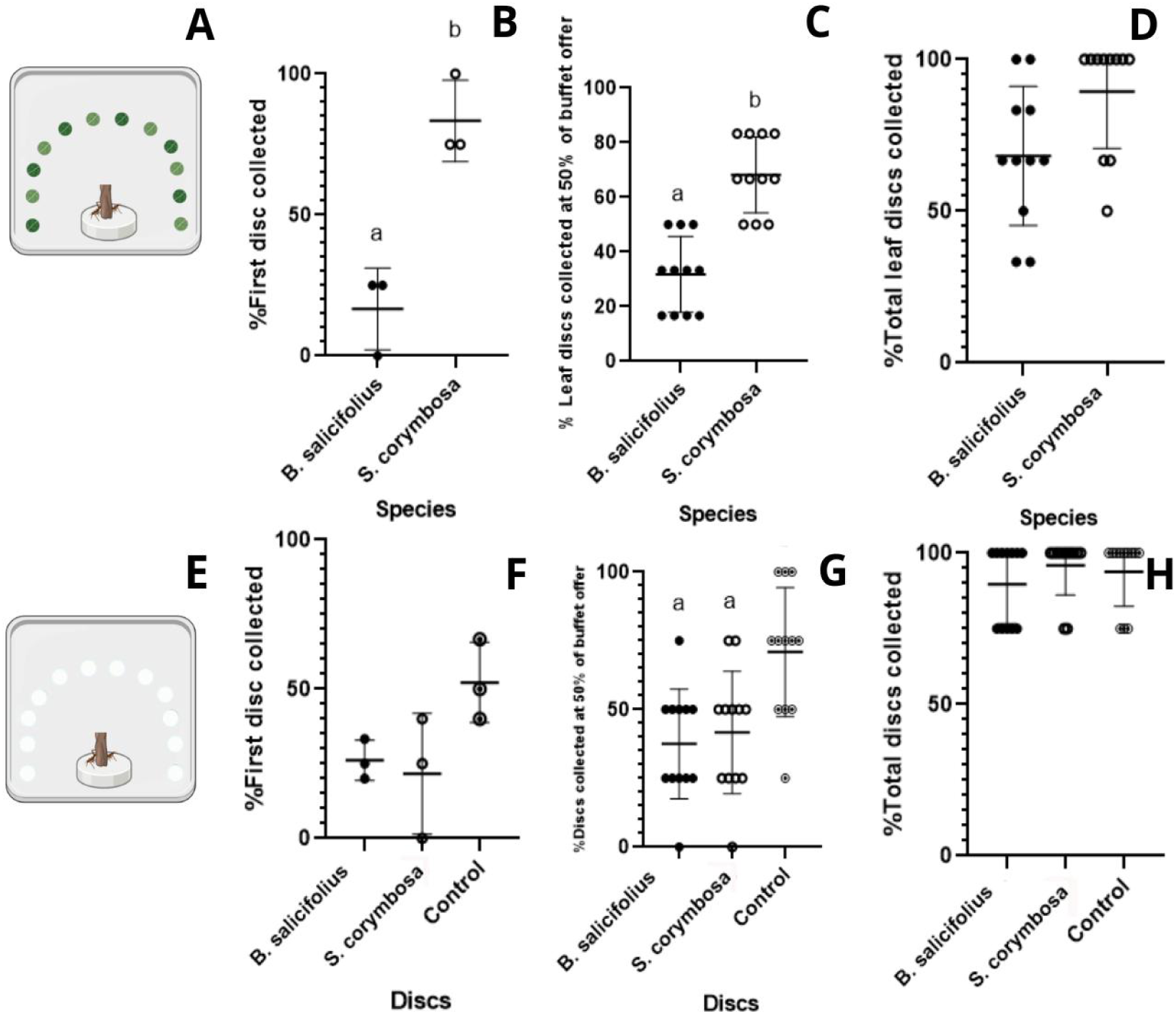
Collective preference assays between least preferred (*B. salicifolius*) and most preferred (*S. corymbosa*) leaf discs. A. Experimental arena layout for selecting fresh leaf discs. The distance from the bridge drop to each disc was 6 cm. B. Percentage of trials in which the first disc collected was *S. corymbosa* or *B. salicifolius*. Each point corresponds to a colony. The mean and standard error are also indicated. C. Percentage of discs collected when 50% of the initial supply was reached. Each point represents the result of one replicate of the experiment. D. Percentage of discs collected at the end of the experiment. Data from 3 colonies, 4, and 3 independent replicates in each. E. Experimental arena to evaluate the role of waxes, which are detected by ants through contact, by the same design as in A. Paper filter discs with extracts of waxes from the plant species under study were used. Data from 3 colonies, between 5 and 3 trials were performed with each one on different days. The three points on each disc type indicated on the X axis correspond to the proportion of replicates in which each colony raised that disc as the first choice. The horizontal bars and error bars indicate the mean and standard deviation of the 3 colonies. F. Idem than B No significant differences were found with respect to a random distribution (χ2=1.51; gl=2; p=0.472). G. Idem to C, different letters indicate significant differences between groups and with respect to a random distribution (X^2^=6.33; gl=2; p=0.042) with a significant preference for control discs (X^2^=5.57; gl=1; p=0.018). H. Percentage of discs with *B. salicifolius*, *S. corymbosa*, or control wax extract, collected at the end of the trial with no significant differences (X^2^= 30.48; gl=22; 0.1072).

To measure the preference of foraging ants for fresh leaf discs of *B. salicifolius* and *S. corymbosa*, the following data was recorded: i) the species of the first disc collected, and ii) the species of the first 6 discs collected. The analysis of the data corresponding to the first disc was based on the proportion of replicates of the experiment with the same colony in which the first disc was *B. salicifolius* or *S. corymbosa*, and therefore a single proportion value was obtained per colony. To analyze the first 6 discs, the proportion of *B. salicifolius* and *S. corymbosa* discs in each trial was taken into account, and therefore the results are expressed as the proportion of each species in each replicate of the experiment. Both constitute alternative and complementary analytical approaches.

To evaluate the role of leaf surface waxes, which are inspected by ants through contact, filter paper discs were used instead of fresh leaf discs, and wax extracts from the plant species under study were added to each of the filter paper discs (Fig. 3.E). The extracts were made by immersing 10g of leaves from each of the native species in 50 ml of dichloromethane (pesticide grade - Sintorgan) for 20 seconds (Cameron et al., 2002). The dichloromethane was then evaporated under a stream of nitrogen until the solution reached 2 ml. Three μl of solution was placed on each disc to be used in the behavioral arena, and the solvent was allowed to evaporate. Each colony was then presented with a group of 12 filter paper discs containing wax extracts from the study species and control discs (solvent only). These groups consisted of four discs containing *B. salicifolius* extract, four discs containing *S. corymbosa* extract, and four control discs to measure spontaneous collection of paper discs. The tests were conducted on three colonies and repeated two to four times for each colony. Selectivity indices were calculated as in the collective preference test with fresh leaf discs.

### Individual preference assays least preferred (*B. salicifolius*) and most preferred (*S. corymbosa*)

Each colony was connected from its own collection arena to a new experimental collection arena (43cm x 11cm x 6cm) by means of a wooden bridge which offered *ad libitum* oat flakes. Once an ant returned with an oat flake, it was picked up with forceps from the same flake and transferred (ant+flake) to the end of a second experimental device. The latter varied between a bridge associated with a platform (Fig. 4.A) and a bridge associated with a stationary ambient olfactometer (Fig. 4.F), according to Saverschek & Roces (2011) methodology. After moving the ant and its flake to the end of the experimental bridge, the oat flake was removed, ensuring minimal disturbance to the ant so that it would naturally continue its foraging behavior. This technique allowed for accurate identification of ants motivated to forage.

**Figure 4.**
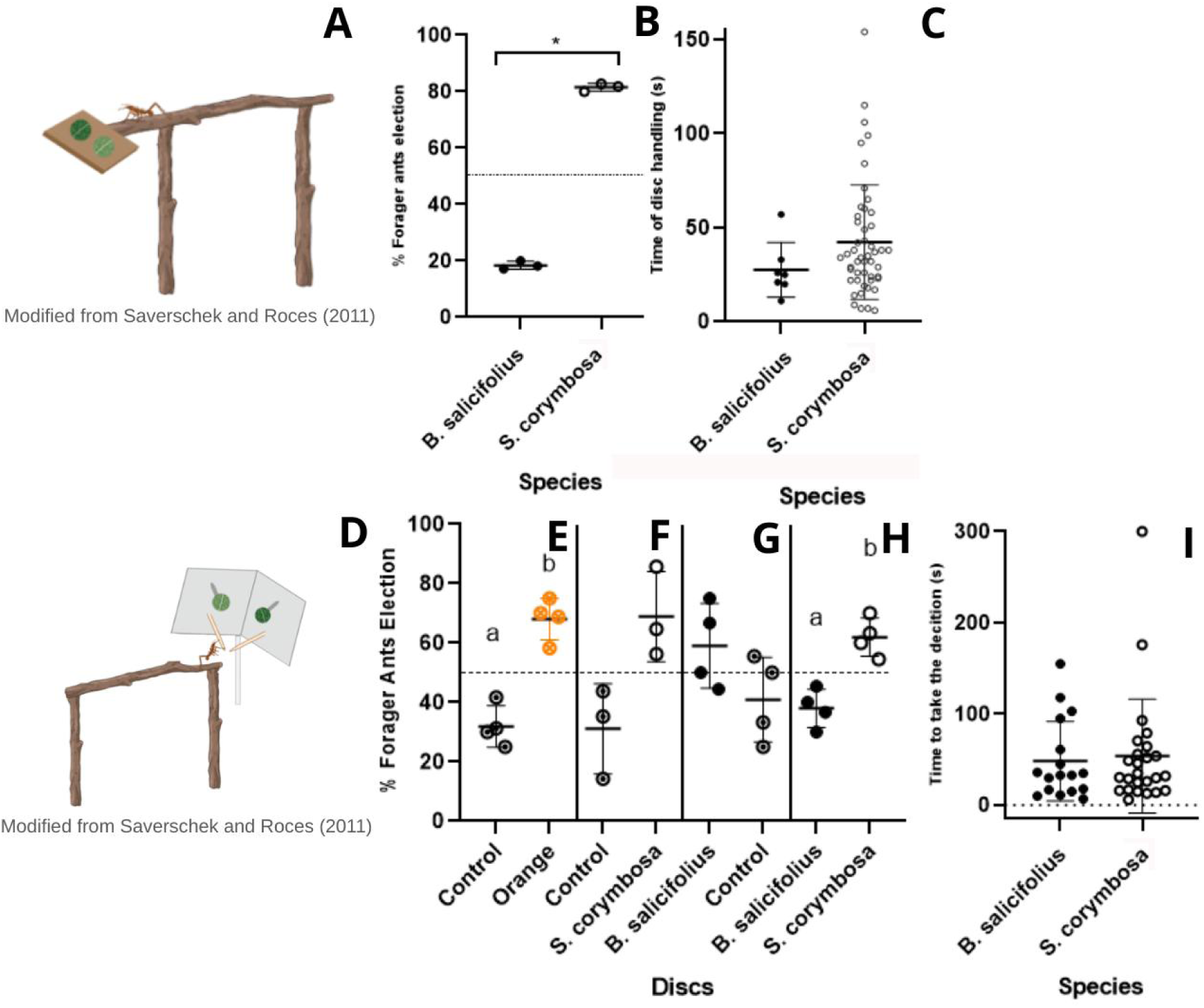
Individual preference assays between least preferred (*B. salicifolius*) and most preferred (*S. corymbosa*) plant species. A. Schematic of individual collection platform with a choice between two discs of fresh *B. salicifolius* and *S. corymbosa* leaves. 3 colonies were used. B. Percentage of disc selection between *B. salicifolius* or *S. corymbosa*. Each point indicates the percentage of times that an individual from a given colony collected *B. salicifolius* or *S. corymbosa* (n=106; X^2^=37.24; gl=1; p<0.01). The graph also shows the mean and standard deviation. C. Time to handle a disc (seconds) until collection; 8 *B. salicifolius* and 51 *S. corymbosa* discs were handled (n=59; t test, nonparametric; p=0.077). D. Schematic of a stationary ambient olfactometer with fresh discs of the native plant species *B. salicifolius* and *S. corymbosa*, adapted from (Saverschek and Roces, 2011). E-H. Percentage of forager ant choices in a stationary ambient olfactometer with a choice between two odor sources. The graphs represent the percentage of decision-making events toward one source or the other for each colony. The mean and standard deviation between colonies are also presented. The odorant sources were: E. Control vs. Orange essence, (n=50; X^2^=6.48; df=1; p=0.01). F. Control vs. *S. corymbosa* discs, (n=80; X^2^=7.2; df=1; p<0.01). G. *B. salicifolius* discs vs Control, (n=38; X^2^=2.63; df=1; p=0.10). H. *B. salicifolius* disc vs. *S. corymbosa* discs (n=92; X^2^=5.26; df=1; p=0.02). I. Time required (seconds) for an ant to make the decision to approach one of the two odor sources coming from *B. salicifolius* a or *S. corymbosa* discs.

### Contact cues: Collection platform

Three colonies of *A. ambiguus* were used, 106 ants responded actively and chose one of the two discs presented. Fresh leaf discs of native plant species, *B. salicifolius* and *S. corymbosa*, were close enough to each other that the ant could make physical contact and integrate olfactory and contact cues with both of them before deciding to pick one up. The discs were placed very close to each other (approximately 0.5 cm), so it is highly unlikely that an ant would pick up a disc simply because it was the first one it found (Fig. 4.A). The ant was placed at the distal end of the bridge. We later recorded the selected plant species and the time it took each ant to handle one of the leaf discs until they picked it up (hereafter referred to as handling time). A foraging event was considered when an ant picked up a disc, placed it on its head and/or thorax, and took a step with it.

### Volatile cues: Olfactometer

To evaluate the individual preference of forager ants for odorants from fresh discs, an stationary ambient olfactometer (Saverschek and Roces, 2011) was used (Fig. 4.D). A positive control consisted of comparing clean filter paper with a second-one containing orange essential oil. For the trials with native plants, four colonies of *A. ambiguus*, and a total of 92 animals were used.

Each experimental ant was placed at one end of an intermediate bridge. During the experiment, the ant was able to move across the bridge to the other end, where the device was placed. This device consisted of a thin cardboard pierced by two wooden sticks placed at 90° angles to each other. A smaller metal needle was placed at the same angle on each of them, with a disc of fresh leaves or filter paper at its end, balanced as shown (Fig. 4.D). The discs of fresh leaves or filter paper were placed out of the ant’s reach, so that, in order to approach the odor source, the ant had to decide which wooden stick to orient itself toward, take a step toward it, and then climb onto it. Therefore, each ant’s decision to move to one stick and the time for it were recorded. After use, each device was discarded except for the metal needles, which were reused after being washed in 96% alcohol. The control sample consists of a filter paper disc treated with 3μl of 1% orange essential oil (in dichloromethane) and a control disc (dichloromethane). The experiment was then conducted with discs of fresh, intermediate-ripe leaves from various specimens of the native *B. salicifolius* and *S. corymbosa* species.

### Leaf Toughness

To determine the toughness of the leaves of the native plant species studied, the force required to penetrate the leaf and the lignin content were analyzed.

### Physical Measurements

A Medio-Line 41002 penetrometer (Pesola, Switzerland) with a tip diameter of 1.9 mm and a circular area of 2.83 mm^2^ was used. From each of the plant species analyzed, 10 leaves of intermediate maturity were randomly selected and crossed perpendicularly until the tip pierced the leaf. Measurements were taken at 10 points located around the leaf perimeter to ensure samples were obtained along the leaf’s length. For each measurement, the penetrometer reading was recorded and divided by the circular area of the penetrometer. Using the 10 data points corresponding to each leaf of the same species, the average per species was obtained. This value is an indicator of leaf hardness.

### Lignin

Lignin was measured through sequential extractions based on fiber analysis (Van Soest, 1963). An Ankom 220 fiber analyzer (Ankom Technology) was used, which includes acid detergent digestion, subsequent sulfuric acid digestion, and ash determination. Four *B. salicifolius* and four *S. corymbosa* samples were analyzed.

### Specialized metabolites and nutrients

#### Volatile Organic Compounds (VOCs)

VOCs were collected from the aerial sections of specimens grown in pots and maintained in the lab. Prior to collection plants underwent a 48-hour acclimatization in the chamber. Volatiles were collected under controlled conditions (25 ± 2°C, L16:O8, light intensity 350μmol/ms). The aerial parts of the plant species were placed inside transparent polyethylene terephthalate (PET) bags (Toppits oven bags, Minden, Germany) that enclosed approximately 1 m of specimen length in an air current circuit that traversed the plant headspace. A push-pull pump (model AT-703, Atman, China) was used to initiate the air current. The filter consisted of a metal tube (19.5 m long, 4.5 cm internal diameter) with granulated activated carbon (120 g). The air then passed through a flow meter (model 112-02-A, Aalborg Instruments, Orangeburg, New York, USA). The volatile collection trap consisted of a Pyrex glass tube (7.5 cm long, 4 mm internal diameter) containing 30 mg of HayeSep Q adsorbent. A suction pump (Standard model, Tuff, Bedford, UK) generates negative pressure, forcing the air inside the PET plastic bag through the collection trap.

Volatiles were collected from four non-damaged *B. salicifolius* and eight *S. corymbosa* specimens, using PET plastic bags without plants as blanks. Each *B. salicifolius* specimen was measured in a separate plastic bag, while two *S. corymbosa* specimens were measured in another plastic bag. This compensates for differences in size and/or foliage (in plants of the same age, *B. salicifolius* has twice the foliage of *S. corymbosa*). Volatile collection lasted 6 hours (from 10:00 a.m. to 4:00 p.m.); the collection traps were wrapped in aluminum foil and stored in a freezer (−20°C) until elution with 150 μl of dichloromethane (Pesticide grade - Sintorgan) containing 5 ng/μl of dodecane (Sigma-Aldrich) as an internal standard. Samples were analyzed with GC-MS, Agilent 7890-A coupled to an Agilent 5977 selective mass detector), with a DB5-MS capillary column (0.25 mm internal diameter x 0.25 μm film thickness). Samples (1 μl) were injected at 240°C in splitless mode. Helium as carrier gas with flow rate of 0.7 ml per minute (inlet pressure: 20.48 kPa). Column temperature at 35°C for 1 minute and then increased at a rate of 10°C per minute until 230°C for 15 min. Data were analyzed using MassHunter Workstation software (version B.06.00, Agilent). Compounds were identified by comparing their mass spectra with those provided by the NIST Chemistry WebBook (William E. Wallace, director, “Mass Spectra”) and “Identification of essential oil components by gas chromatography/mass spectrometry” book (Adams, 2007). Results were compared with the NIST database (National Institute of Standard and Technology, USA). Identification by Kovats index (KI, Eq. 1) was corroborated compared with tabulated values for a DB5-MS column published by NIST (William E. Wallace, director, “Retention Indices”) and/or by Adams (2007). The Kovats index of a component in a sample is an indicator of the relationship between the retention time of a component in the sample and the retention times of two successive alkanes in a standard series of alkanes eluting immediately before and after the component of interest (Ettre, 1993). Only identified compounds that were found in at least 3 of the 4 replicates are shown.

The Kovats indices of the analyzed compounds were calculated using “The Pherobase” (El-Sayed AM 2022. The Pherobase: Database of Pheromones and Semiochemicals. https://www.pherobase.com).

### Extraction of Total Polyphenols

Processed plant material (40–80 mg) was extracted and 50% methanol solution was added at 80°C in a heat bath for one hour. The extract was then filtered (Whatman No. 1). A 0.5 ml aliquot was added to a colorimetric solution consisting of 10 ml of distilled water, 1.25 ml of the Folin-Ciocalteu reagent (Cadish and Giller, 1997), and 5 ml of 17% sodium carbonate, in that order, diluted with distilled water in 25 ml. Absorbance was read (λ=760nm) with Spectronic Genesys 2 spectrophotometer (Spectronic Instruments®). For quantification, a calibration curve was made, and gallic acid (0.1 mg/ml), was used as a standard according to Vivanco and Austin (2008) protocol.

### Total Sugars

Dubois et al. (1956) protocol was followed with 0.035 g of plant material using the phenol-sulfur method: 7 ml of 2.5 N HCl, incubated at 100°C for 3 hours. The solution was then neutralized with sodium carbonate (NaCO3), brought to a volume of 100 ml. One ml of the above extract was taken and 1 ml of 5% phenol and 5 ml of 96% concentrated sulfuric acid were added. After 10 min, the solution was heated to a 25-30°C temperature for 20 min. Absorbance was read (λ=490 nm). Values obtained were compared with a calibration curve made with 100 mg/L glucose standard in distilled water. Seven samples of *B. salicifolius* and seven samples of Sen from the field were used, each from a different specimen.

### Organic Matter

The sample mass was weighed before and after being transferred to the furnace. Ash content was subtracted from the initial weight as a measure of organic matter loss (Harmon et al. 1999). One crucible per sample and the sample to be analyzed (0.20-0.28 g) were weighed. Samples were placed in a muffle furnace at 400°C for 4 hours. Each sample was reweighed to infer the amount of organic matter. Known amounts of fescue (*Lolium arundinaceum*) were used as an organic matter standard. Seven samples from *B. salicifolius* and seven samples from *S. corymbosa* were used.

### Total Proteins

Bradford soluble protein measurement protocol was adapted. Cys PIs extraction buffer (50 mM NaPO4, 150 mM NaCl, 2 mM EDTA, pH 7.2) was added in a 1:3 ratio (10 mg sample with 300 µl of buffer). The sample was homogenized by vortexing and centrifuged for 15 min at 4°C and 12,000 rpm. Supernatant was removed and centrifuged in equal conditions. Each sample was quantified in a microplate reader (Varioskan Lux Multimode Microplate Reader by ThermoFisher) with 10 µl in 200 µl of Bradford standard (BioRad), previously prepared with distilled water in a 1:4 ratio. The reader was shaken for 1 min, and the absorbance was read (595 nm). The calibration curve was run by duplicate, with 6 points between 0 and 1 µg/ml of BSA in distilled water. Eight samples from *B. salicifolius* and six samples from Sen from the field were used; all were measured by duplicate.

### Statistical Analysis

Statistical analyses were performed using Graphpad Prism 8.0.1 software for Windows. The t test and chi-square test were used. One-way ANOVA with Duncan’s post hoc comparisons were also used to compare multiple treatments when the data distribution was normal and had homogeneity of variance. The Kruskal-Wallis test and Dunn’s post hoc comparisons were used when the data distribution was not normal. The analyses are described in more detail in each experimental section as appropriate.

### AI

AI tools were used to improve human-generated texts for readability and style, and to ensure that the texts are free of grammar errors.

## Results

### Herbivory measurements

The total foliage of the plant species selected for evaluation was measured in three native plant patches. The three patches presented initially different foliage amounts, distributions, compositions, light intensity, size, and number of specimens evaluated. This variability reflects the natural heterogeneity of the environment in which the ants normally forage. Despite this diversity, all patches could be used for measurements throughout the summer and fall months (Fig. 1). The presence of foliage was systematically quantified for *B. salicifolius, M. laetevirens*, *E. crista-galli* and *S. corymbosa*, (N=4). While it cannot be confirmed that ant activity was the sole cause of foliage loss, the type of damage and consumption were consistent with leaf-cutting ant damage. Figure 1 shows the loss of foliage in *S. corymbosa* specimens during the season, while the remaining species did not show this level of foliage loss. Importantly, these field observations provided an important basis for selecting plant species and designing the subsequent quantitative laboratory experiments.

### Consumption hierarchy in the laboratory

We established a hierarchy of consumption preferences in *A. ambiguus* based on pairwise presentations of leaves from eight native plant species across six colonies. Significant differences in consumption were observed among plant species (one-way ANOVA; F=10.40; df=7,40; p<0.001), allowing the identification of species that were consistently more or less consumed than others (Fig. 2). Species occupying adjacent positions in the hierarchy did not differ significantly from one another. In contrast, significant differences (p<0.05) were found between each species and the non-adjacent species in the hierarchy (Fig. 2.C.3). Therefore, species located lower in the hierarchy were less preferred than those ranked higher. Among the least preferred species, *B. salicifolius* and *M. laetevirens* stood out, whereas *S. corymbosa* and *V. bonariensis* were among the most preferred. Figure 2.E shows the same data separated by colony. As can be observed, preferences varied among colonies, although all colonies showed similar overall trends, particularly regarding the most and least preferred plant species (Fig. 2E).

Consumption trials were video-recorded in five of the six colonies and subsequently analyzed to gain further insight into choice and preference behaviors. This approach allowed us to divide consumption behavior into the following stages: (i) approach and first contact, (ii) first cut, and (iii) consumption (see Materials and Methods). For each stage, the eight plant species were ranked using a methodology similar to that applied for overall consumption. Interestingly, we repeatedly observed that some plant species that initially elicited strong attraction were ultimately among the least consumed. Figure 2.D shows the plant preference hierarchies across the three behavioral stages. Final consumption could not be predicted from the initial approach, but was strongly associated with the first cut stage. We evaluated the percentage of coincidence between the ant’s first approach and the first cut, as well as the percentage of coincidence between the first cut made on a leaf and final consumption. The 28 trials conducted with each colony were used for this analysis, and the percentage of coincidence between the 28 “approaches”, the 28 “first cuts” was calculated and similarly for the 28 “first cuts” with the 28 “final consumptions”. A statistical analysis based on Chi-square comparisons against a random distribution showed significant differences only for the second stage, p<0.001 (Fig. 2.F).

From these results, we can conclude that a first approach is not a good predictor of final consumption of the fresh leaf. However, the first cut is. Therefore, it is highly important to analyze what types of sensory cues are involved in the initial approach and which ones are involved from contact to the first cut, leading to the consumption of the plant material. Consequently, in the next sections we studied the relevance of different physicochemical cues, as well as collective and individual behaviors that may guide plant consumption.

Collective preference assays between least preferred (*B. salicifolius*) and most preferred (*S. corymbosa*)

To provide a more comprehensive description of foraging preferences and their relationship to different types of sensory cues, we focused the next experiments on two plant species that showed clearly different preference profiles and that, due to their availability and general characteristics, could be used in reforestation projects and push-pull ant management programs. Therefore, we chose *B. salicifolius* and *S. corymbosa* as extreme examples within the consumption preference hierarchy.

For a collective foraging experiment we use a foraging assay of 12 fresh leaf discs —6 from *B. salicifolius* and 6 from *S. corymbosa*— arranged in a collection arena. Three ant colonies were evaluated, and between 3 and 4 measurements were taken per colony, each on different days (a total of 11 replicates). The percentage of replicates in which the first disc collected was either *S. corymbosa* or *B. salicifolius* was determined for each of the three colonies (Fig. 3.B). To assess whether the collection of the first disc differed from a random distribution (50/50), an exact binomial test was performed. The results indicated a significant deviation from the expected distribution (X^2^=4.45; p=0.035). To determine whether this collection pattern for *S. corymbosa* and *B. salicifolius* depended on the colony, a Chi-square test of independence was performed, which indicated no differences between colonies (X^2^=0.92; p=0.63). Therefore, we observed a clear preference for *S. corymbosa* in regarding the first collected disc.

The progression of collection was then evaluated, taking into account the percentage of *B. salicifolius* and *S. corymbosa* discs collected when collection reached 50% of the initial supply. Fig. 3.C shows the percentage of each species collected at that time for each replicate of the experiment. Significant differences were found between colonies or replicates, and the number of discs collected from *S. corymbosa* and *B. salicifolius* was found to differ from a random distribution (X^2^=8.73; gl=1; p=0.0031). Therefore, at 50% of the initial offer, there is a difference in the preference between the two species. Fig. 3.D shows, for each replicate of the experiment, the final percentage of *S. corymbosa* and *B. salicifolius* discs collected at the end of the maximum experiment time. The observed median was 69.45 (*B. salicifolius*) and 91.68 (*S. corymbosa*), and no significant statistical differences were found (Wilcoxon; p=0.1250).

In summary, we found that *S. corymbosa* discs were collected more quickly by the ants, so we interpret them as preferred over those from *B. salicifolius*. However, the fact that the *B. salicifolius* discs were also collected, albeit at a slower rate, suggests that this is not a rejection of the discs, but simply a lower preference.

### Role of cuticular waxes

We studied the contribution to ant preference for *S. corymbosa* and *B. salicifolius* of cuticular waxes, since they are one of the physical defense mechanisms of plants (Erb and Reymond, 2019) as well as a possible source of recognition cues. To this end, we analyzed each colony’s preference for 12 filter paper discs containing the cuticular wax extracts. The experimental arena contained 4 discs with *B. salicifolius* cuticular wax extracts, 4 discs with *S. corymbosa* cuticular wax extracts, and 4 paper discs containing solvent as a control. The experiment was conducted with 3 ant colonies, and between 5 and 3 trials were performed per colony (12 complete measurements).

For each colony, the percentage of times that the first collected disc contained extracts of cuticular waxes from *B. salicifolius*, *S. corymbosa*, or neither was analyzed (Fig. 3.F). The distribution of the first collected discs was analyzed without finding significant differences with respect to a random distribution (X^2^=1.51; gl=2; p=0.472). Therefore, there are no preferences for any of these discs in the first collection made (Fig. 3.F). As an alternative measurement of a differential preference between discs, the collection profile of the first 6 discs of each trial was analyzed. We observed that the percentage of each type of disc collected up to 6, that is, 50% of the offer, shows no significant differences with respect to a random distribution between the native species (X^2^=5.57; gl=1; p=0.018) while comparing the number of discs in each category versus a random collection of discs showed significant differences only for the group of discs without cuticular wax extracts and significant differences with respect to a random distribution between native species and control discs (X^2^=6.33; gl=2; p=0.042), (Fig. 3.G). Therefore, there is a preference for discs without cuticular extracts, the control group, while this preference becomes less evident as these discs are less represented in the experimental arena. Finally, the total number of discs collected was recorded when the test time was completed (Fig. 3.H) and it was verified that practically 100% of the discs were collected regardless of the origin and the presence or absence of cuticular wax extracts showing no significant differences between them (X^2^=30.48; gl=22; p=0.1072).

Based on the experiment and the analysis carried out, it can be concluded that the cuticular wax extracts from *S. corymbosa* and *B. salicifolius* do not play a determining role in the preference indices between both species. Based on the observed trends (Fig. 3) and differences in collection at 50% of discs (Fig. 3.G), it may be speculated that the cuticular wax extracts provoke rejection rather than acceptance, which is more noticeable in the case of *B. salicifolius*. This last interpretation arises from observing that the ants also collected control discs without cuticular wax extracts and that the trends suggest that they are more preferred than those treated with cuticular wax extracts from *S. corymbosa* and *B. salicifolius* (Fig. 3).

### Individual preference assays between least preferred (*B. salicifolius*) and most preferred (*S. corymbosa*) plant species

Next, we studied whether the preferences observed in the previous section are maintained at the individual level. Therefore, we worked with two types of devices that allowed us to evaluate different types of sensory cues and individual decisions.

### Contact cues: Individual Collection Platform

To evaluate the role of contact cues in ant preference, we used a small foraging arena in which ants could simultaneously contact two different leaf discs, ensuring that the foraging decision was made only after contacting both options. Under these conditions we observed a clear preference to collect *S. corymbosa* discs over those from *B. salicifolius* (Fig. 4.B). The observed distribution differed significantly from the distribution expected under random disc collection (n=106 from 3 colonies; X^2^=37.24; df=1; p<0.01). As shown in Fig. 4.C, no significant differences were found in the handling time of the two types of discs (n=59; t test, nonparametric; p=0.077). These results suggest that contact cues from *B. salicifolius* are not repellent to the ants; rather, *B. salicifolius* is simply less preferred than *S. corymbosa*.

### Volatile cues: Stationary Ambient Olfactometer

The next experiment was performed to evaluate exclusively the role of olfactory cues. For that aim we used a stationary olfactometer in which ants decide to approach one of two plants based only on volatile perception. We started doing a positive control experiment using orange essence. The percentage of individuals in each colony that oriented toward the orange side or the clean filter paper disc was analyzed, revealing a significantly greater preference for orange (Fig. 4.E). This pattern was homogeneous across colonies and significantly different from random choice (n=50 from 4 colonies; X^2^=6.48; df=1; p=0.01). Following a similar experimental design we tested the attractivity of *S. corymbosa* volatiles. Following a similar experimental design, we then tested the attractiveness of *S. corymbosa* volatiles. A significantly greater number of individuals approached the *S. corymbosa* discs than expected under a random distribution, indicating attraction to *S. corymbosa* volatiles (n=80 from 4 colonies; χ²=7.2; df=1; p<0.01) (Fig. 4.F).

When comparing *B. salicifolius* discs with control discs, we observed a tendency for a greater number of individuals to approach the *B. salicifolius* disc. However, the distribution of choices did not differ significantly from random expectation (n=38 from 4 colonies; χ²=2.63; df=1; p=0.10) (Fig. 4.G).

Finally, we tested *B. salicifolius* against *S. corymbosa*. The preference was evaluated using 92 individuals from four colonies. We found a significantly greater number of individuals approaching *S. corymbosa* discs than expected under a random distribution, indicating a preference for *S. corymbosa* over *B. salicifolius* (n=92 from 4 colonies; χ²=5.26; df=1; p=0.02) (Fig. 4.J). No differences were found in the time required to make decisions toward *S. corymbosa* or *B. salicifolius* discs (Fig. 4.K).

### Specialized metabolites and nutrients

#### Volatile Compounds: Collection and Analysis of Volatile Organic Compounds

Chromatographic profiles were obtained for *B. salicifolius* (Fig. 5.A) and *S. corymbosa* (Fig. 5.B) volatile compounds. Samples from both species were standardized to an equivalent amount of foliage prior to volatile collection. The species presented qualitative and quantitative differences in the emission of volatile compounds. Eleven volatile organic compounds were identified for *B. salicifolius* and seven for *S. corymbosa* (Fig. 5.C-D). Only one volatile compound was common to both species, α-Pinene, although only traces were found in the case of *S. corymbosa*. The average of total volatiles from *B. salicifolius* were 84.23 ng/µl, while those from *S. corymbosa* were 0.44 ng/µl, relative to the internal standard in both cases.

**Figure 5.**
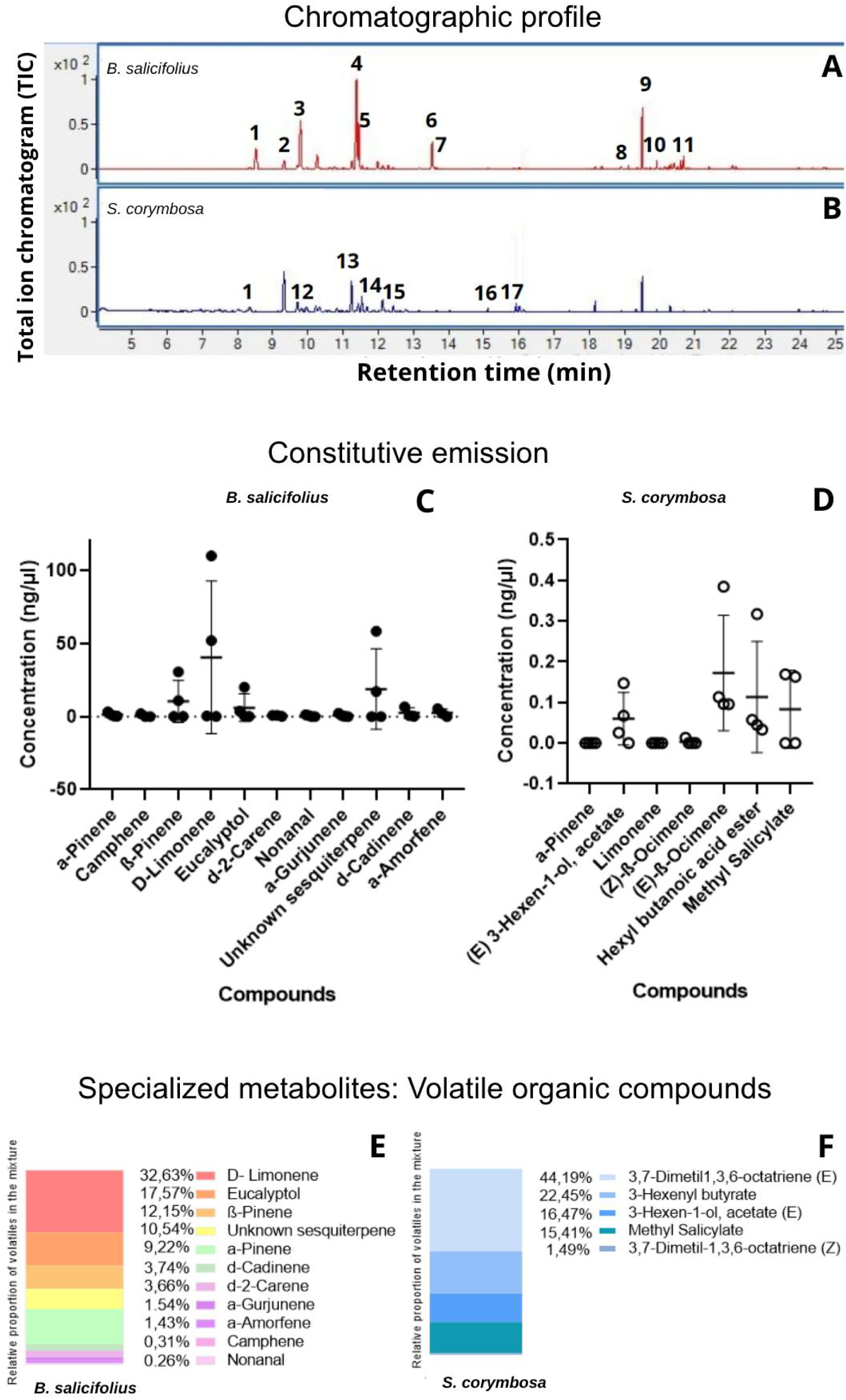
Collection and analysis of volatile organic compounds. A and B. Chromatographic profiles for *B. salicifolius* and *S. corymbosa*. The y-axis represents the total ion chromatogram in arbitrary units (TIC) as a function of the retention time of each compound, measured in minutes. The numbers above the peaks indicate the compounds identified and listed in Supplementary Table 1 and included in Fig. 5.E-F. C and D. Constitutive emission of volatile organic compounds from *B. salicifolius* and *S. corymbosa*. Means ± standard errors are shown. Biological replicates N=4 (where each replicate contains one plant of *B. salicifolius* and two plants of *S. corymbosa* to equalize for foliage). E and F. Proportion of volatile compounds found in *B. salicifolius* and *S. corymbosa*.

The two species differ not only in the absolute concentration of volatiles, but also in their chemical compositions. *B. salicifolius* volatile blend is dominated by approximately 75% monoterpenes, particularly limonene and eucalyptol, and 17% sesquiterpenes, dominated by a non-identified compound that presents similarities to α-cubebene (Fig. 5.E). In the case of *S. corymbosa*, the blend is composed of approximately 45% monoterpenes (mainly ocimene) and 38% green leaf volatiles, particularly esters derived from 6-carbon chains. In addition, it consists of approximately 17% methyl salicylate, a methyl ester derived from salicylic acid (Fig. 5.F).

#### Physicochemical Measurements

Different physicochemical analyses were made for both species (Table 1). *B. salicifolius* presented significantly higher leaf toughness (t test, p=0.0036), higher amount of total polyphenols (t test; p<0.0001) and more organic matter (t test; p=0.0015). Meanwhile *S. corymbosa* presented significantly higher levels of lignin (t test; p=0.0006). Both species show no significant differences in total sugars (t test; p=0.3074) and a trend, although not significant, was observed towards a higher amount of total proteins in *S. corymbosa* (t test; p=0.0749) (Table 1).

**Table 1.** Physicochemical characteristics of least preferred (*B. salicifolius*) and most preferred (*S. corymbosa*) species were analyzed. Leaf toughness along its perimeter (N=10 leaves, n=10 measurements per leaf). Percentage of lignin content (N=4). Total polyphenols (N=7). Total sugars (N=7). Organic matter (N=7). Total proteins (*B. salicifolius* N=7, *S. corymbosa* N=6). For all physicochemical characteristics t tests were made.

| Analyzed characteristics | *B. salicifolius* | *S. corymbosa* | p value |
| --- | --- | --- | --- |
| Force (g/mm<sup>2</sup>) | 198.25 ± 4.91 | 167.3 ± 3.68 | 0.0036 |
| Lignin content (%) | 4.24 ± 1.20 | 9.82 ± 0.41 | 0.0006 |
| Total polyphenols (mg/g) | 0.37 ± 0.026 | 0.05 ± 0.0040 | <0.0001 |
| Total sugars (mg/g) | 539.3 ± 26.20 | 514.0 ± 31.70 | 0.3074 |
| Organic Matter (%) | 0.920 ± 0.0040 | 0.900 ± 0.0050 | 0.0015 |
| Total proteins (mg/ml) | 0.180 ± 0.038 | 0.480 ± 0.033 | 0.0749 |
| Relation between total proteins & total sugar | (3.33 ± 0.72) $\times 10^{-4}$ | (9.33 ± 0.86) $\times 10^{-4}$ | - |

When analyzing the relationship between total protein and total sugars (Table 1), we found that *S. corymbosa* values greatly exceed those of *B. salicifolius*. Therefore, the relationship between total proteins and total sugars could be an indicator of the nutritional value of these species relevant to harvester ants.

## Discussion

Our results provide novel ecological information in relation to native flora of the Lower Parana River Delta and leaf-cutting ant herbivory. Integration of these native species into *Salix forest plantation* management could simultaneously enhance pest control and contribute to ecosystem restoration, with cascading benefits for associated fauna, including insectivorous birds. Specifically, *V. bonariensis* was consumed significantly more than the commercial willow species *S. babylonica*, which, due to its herbaceous and perennial nature, makes it a prominent candidate for push-pull strategies (Perri et al., 2020), since if cultivated artificially it could act as alternative vegetation for the ants. Meanwhile, *S. corymbosa* appeared to be a strong attractant, although its woody, shrub-like growth habit could make its cultivation and maintenance more challenging in practical applications. Conversely, *M. laetevirens*, *B. salicifolius*, *E. crista-galli* and *T. australis* were consumed significantly less than *Salix*, suggesting that this native species would be unsuitable as alternative vegetation near willow plantations, as leaf-cutting ants preferentially foraged on the commercial species rather on this native plant.

By integrating field herbivory observations (Fig. 1) with controlled laboratory assays of foraging preference (Fig. 2–4), we consistently identified a robust preference in leaf-cutting ants, where *S. corymbosa* represented a highly preferred resource and *B. salicifolius* a significantly less consumed species. This is specifically relevant since leaf-cutting ants are highly generalist herbivorous, they can forage up to 50% of the available plant species in a plant community (Wirth et al., 2003).

Despite this general trend, consumption hierarchies exhibited colony-specific variation (Fig. 3). These results are consistent with previous reports on colony-specific variability across seasons and years, as well as due to environmental conditions, colony ontogeny, nutritional demands and physicochemical plant traits (Howard, 1987; Meyer et al., 2006; Herz et al., 2007; Montoya-Lerma et al., 2012; dos Santos et al., 2013; Arenas and Roces, 2017; Perri et al., 2020). Nevertheless, the preferred and non-preferred groups across colonies (e.g., *V. bonariensis*/*S. corymbosa* vs. *B. salicifolius*/*M. laetevirens*) suggest common underlying determinants of preference that could be conserved due to sensory and nutritional mechanisms. Hence, in terms of preference, the commercial species evaluated (*S. babylonica*) occupies an intermediate position among the analyzed native species.

Mechanistically, our behavioral analyses demonstrate that resource preference is a multi-stage, sensory-driven process. The first cutting event reliably predicted the final consumption of the resource, when ants were presented with multiple plant options (Fig. 3.D). Thus, these results indicate that preference occurs early and it is neither random nor governed by resource availability. This supports prior works regarding the key role of olfactory and gustatory cues in foraging decisions (Littledyke & Cherrett, 1976; Roces, 1994). Precisely, olfactometer assays revealed that plant volatile cues alone are sufficient to elicit oriented approach behavior towards the preferred species (Fig. 4.H). Nevertheless, the absence of a clear avoidance behavior indicates the olfactory cues modulate levels of preference rather than a binary preference-rejection response.

Additionally, the magnitude of preference increased when contact cues were available (Fig. 4.B). Meanwhile, the application of cuticular wax extracts reduced item collection (Fig. 3.F). Hence, behavioral responses to volatiles are further modulated by gustatory and mechanosensory cues. Therefore, multimodal integration likely reflects sequential evaluation of plant sensory cues, where long-range detection via olfaction is refined through direct contact with the item followed by its hardness and palatability. The observed reduction in *B. salicifolius* collection when the forager ant could have access to item contact cues supports the hypothesis that these cues amplify discrimination between resources (Fig. 3.F). This is in line with the relevance of different cues modalities in decision making (Kulahci et al. 2008). At the colony level, these preferences are affected by social feedback mechanisms which can either amplify or modulate individual choices. In many of our observations, ants interrupted or were deterred from collection attempts. This behaviour may reflect delayed rejection (Herz et al., 2008) or waste-mediated information transference (Arenas & Roces, 2017). These findings also align with contemporary conceptualization of eusocial colonies as more than superorganisms (Wheeler, 1911), but as self-organized networks which exhibit “inter-identity”, where individual and collective dynamics co-determine behavior (Canciani et al., 2019).

From a functional perspective, colony-level preference must ultimately be interpreted within the framework of the needs of the obligate fungal symbiont. Ants avoid substrates detrimental to the fungus and regulate intake through collective mechanisms (Arenas & Roces, 2016a&b). In this context, the consistency in individual and collective preference for *S. corymbosa* suggests it does not impair fungal cultivation.

Physicochemical analysis revealed that the preferred species (*S. corymbosa*) has lower leaf toughness despite higher lignin content, suggesting compensatory reductions in other structural components affecting leaf hardness. While having a lower concentration of polyphenols and higher protein content may enhance palatability and nutritional value, especially the relation between total proteins and total sugar. In contrast, the non-preferred species (*B. salicifolius*) exhibit greater toughness, higher polyphenol levels, lower protein content and lower relation between total proteins and total sugar (Table 1), consistent with its lower preference in behavioral assays. These findings are consistent with the hypothesis that resource preference is based on a balance between attractive and deterrent compounds probably detected by gustatory sensilla (Howard, 1988; Hertz et al., 2008; Infante-Rodriguez et al., 2020; Smith et al., 2023).

Volatile profiles reveal remarkable differences between *S. corymbosa* and *B. salicifolius*. *S. corymbosa* is characterized by a low concentration blend dominated by Ocimene and green leaf volatiles, whereas *B. salicifolius* volatile concentration is three orders of magnitude higher, dominated by monoterpenes such as Limonene and Eucalyptol (Fig. 5). These differences likely facilitate long-distance discrimination which is consistent with our behavioral findings in the olfactometer assays. Limonene and Eucalyptol are known to possess insecticidal and antimicrobial properties (Hebeish et al. 2008; Skalicka-Wozniak et al. 2009; Oliveira et al. 2010b; Merghni et al. 2023), which may contribute to reduced leaf attractiveness.

In nature, typical odors consist of mixtures of multiple compounds of varying composition and concentrations. Blend perception occurs within a background context that might enhance its recognition (Chan et al., 2018, Jernigan et al., 2020). This framework may explain how *S. corymbosa*, despite its lower volatile emission, remains detectable and attractive due to the ecological relevance of its odor blend. On the other hand, the higher volatile concentration observed in the least preferred species, B. salicifolius, may be explained by a higher density of leaf trichomes, which increase mechanical resistance and might contribute to volatile emission. Thus, trichoma may therefore contribute to reduced foraging preference (Howard, 1988).

Thereby, our research demonstrates that leaf-cutting ants (*A. ambiguus*) foraging decisions emerge from multimodal sensory integration, physicochemical constraints and colony dynamics that ultimately are modulated by the requirements of the obligate symbiotic fungus. This framework advances our understanding of plant–insect interactions while providing a robust foundation for ecologically informed pest management strategies based on restoration of native flora and fauna.

## Acknowledgements.

Authors express their gratitude to: Hugo Chludil for orange essential oil; Guillermina Kubaczka for Bradford standard; Amy Austin for Ankom and chemical analysis reactive; Daiana Perri, Natalia Fracassi, Edgardo Casaubón, Guillermo Madoz, Hugo Rossi, Djoney Gomez, and Agustín Álvarez Costa for field assistance; to Edgardo Casaubón, Vivero Comunitario Ciudad Universitaria (VICCU) and Vivero Chicos Naturalistas for native plants specimens.

## Ethics

This article does not contain any studies with human participants performed by any of the authors. All applicable international, national, and institutional guidelines for the care and use of animals (insects) were followed.

## Author’s contributions

A.N., P.C.F. and F.F.L. designed the study; A.N. performed the experiments, A.N and M.S.M performed chemical analysis, A.N drafted the text and figures A.N., M.S.M, P.C.F and F.F.L elaborated the final version.

