## Supplementary for "Sequential sensory integration drives foraging decisions in leaf-cutting ants: Volatiles, contact cues and phytochemistry"

**Supplementary information**

**Supplementary 1.** . A Control choice experiment was conducted in which two leaf discs from the same plant species were presented simultaneously in order to estimate the expected variability in foraging decisions when both options were identical. The experiment was repeated for each one of the 8 plant species and for 6 ant colonies.

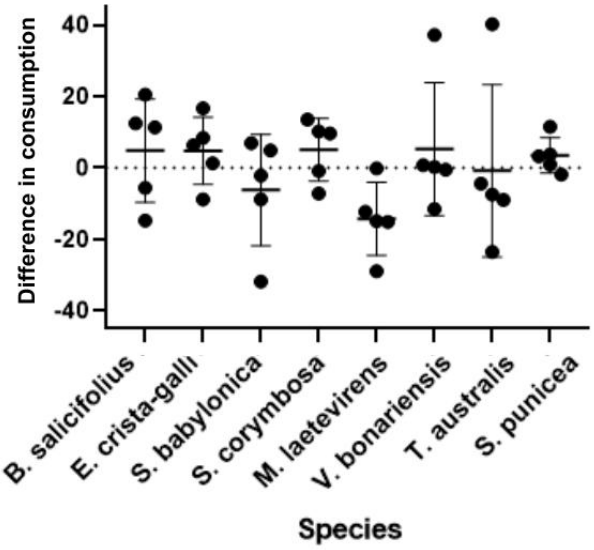


**Supplementary 2. Positive control.** Collection of filter paper discs with cuticular wax extracts from: A. Australian brush cherry (*Eugenia* sp.) or B. Ash (*Fraxinus* sp. ). Four colonies were used and two repeats of the experiment were performed with each colony.

A. Percentage of times that the first disc collected belonged to A or B. Each point corresponds to a colony ($X^2$=2; gl=1; p=0.15). B. Percentage of A or B discs collected when the assay reached 50% (6 discs) of the initial offer. Each point corresponds to a repetition of the experiment ($X^2$=10.08; gl=1; p<0.01) homogeneous distribution between the three colonies ($X^2$=0.096; gl=2; p=0.95).

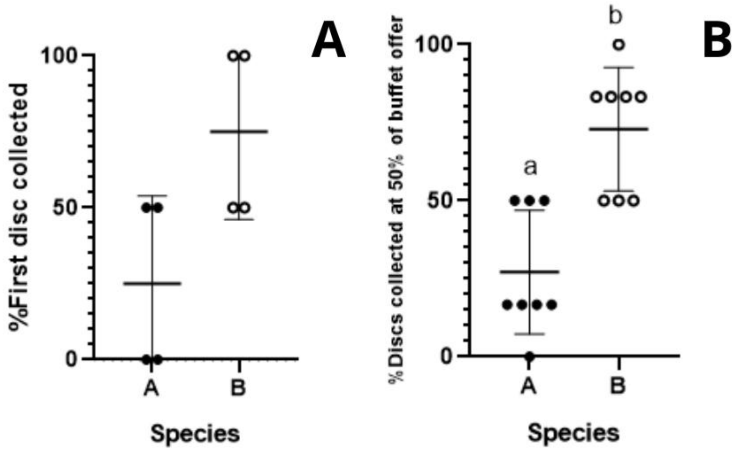

**Supplementary Table 1. Volatile organic compounds identified in *B. salicifolius*. and *S. corymbosa* samples.** For each species, the compound number, the name by which it was identified, the retention time (RT), and the concentration obtained (ng/ul) relative to the internal standard are detailed. Compounds that were not detected in one of the species are symbolized by (-), and those in which only traces were detected are symbolized by (TZ).

| **Compound Nº** | | **1** | **2** | **3** | **4** | **5** | **6** | **7** | **8** | **9** | **10** | **11** | **12** | **13** | **14** | **15** | **16** | **17** |
|---|---|---|---|---|---|---|---|---|---|---|---|---|---|---|---|---|---|---|
| **Compound name** | | α-Pinene | Camphene | β-Pinene | D-Limonene | Eucalyptol | δ-2-Carene | Nonanal | α-Gurjunene | Unknown sesquiterpene | δ-Cadinene | α-Amorphene | 3-Hexen-1-ol, acetato (E) | Limonene | 1,3,6-Octatriene, 3,7-dimethyl- | 1,3,6-Octatriene, 3,7-dimethyl | Butanoic hexenyl ester acid | Methyl salicylate |
| **Retention time (RT)** | | 8,5 | 8,9 | 9,4 | 11,4 | 11,5 | 13,2 | 13,7 | 19,4 | 19,6 | 20,5 | 20,6 | 8,5 | 10,7 | 11,4 | 11,9 | 15,8 | 15,9 |
| **Species** | ***B. salicifolius*** | 1,3 | 0,6 | 10,4 | 40,5 | 6,03 | 0,5 | 0,4 | 0,7 | 18,8 | 2,5 | 2,5 | - | - | - | - | - | - |
| | ***S. corymbosa*** | TZ | - | - | - | - | - | - | - | - | - | - | 0,06 | TZ | 0,003 | 0,2 | 0,1 | 0,08 |
